# iOmicsPASS: a novel method for integration of multi-omics data over biological networks and discovery of predictive subnetworks

**DOI:** 10.1101/374520

**Authors:** Hiromi W.L. Koh, Damian Fermin, Kwok Pui Choi, Rob Ewing, Hyungwon Choi

**Affiliations:** Saw Swee Hock School of Public Health, National University of Singapore, Singapore; Department of Medicine, Yong Loo Lin School of Medicine, National University of Singapore; University of Michigan Medical School, Ann Arbor, MI, USA; Department of Statistics and Applied Probability, National University of Singapore, Singapore; School of Biological Sciences, University of Southampton, Southampton, United Kingdom; Institute of Molecular and Cell Biology, Agency for Science, Technology and Research, Singapore

**Keywords:** integrative-omics, predictive analysis, network signatures

## Abstract

We developed iOmicsPASS, an intuitive method for network-based multi-omics data integration and detection of biological subnetworks for phenotype prediction. The method converts abundance measurements into co-expression scores of biological networks and uses a powerful phenotype prediction method adapted for network-wise analysis. Simulation studies show that the proposed data integration approach considerably improves the quality of predictions. We illustrate iOmicsPASS through the integration of quantitative multi-omics data using transcription factor regulatory network and protein-protein interaction network for cancer subtype prediction. Our analysis of breast cancer data identifies network signatures surrounding established markers of molecular subtypes. The analysis of colorectal cancer data highlights a protein interactome surrounding key proto-oncogenes as predictive features of subtypes, rendering them more biologically interpretable than the approaches integrating data without *a priori* relational information. However, the results indicate that current molecular subtyping is overly dependent on transcriptomic data and crude integrative analysis fails to account for molecular heterogeneity in other -omics data. The analysis also suggest that tumor subtypes are not mutually exclusive and future subtyping should therefore consider multiplicity in assignments.

Availability: https://github.com/cssblab/iOmicsPASS

## Introduction

Today’s systems biology research frequently employs a combination of -omics platforms, seeking to identify systemic biological signals based on comprehensive molecular measurements. Technological advances of the past decades, especially in massively parallel sequencing and mass spectrometry, have propelled a researcher’s ability to profile different areas of “-omics” from the same biological samples. However, interpreting the results from such a large-scale data collection is challenging, and it requires proper computational tools to summarize the relationship between different pieces of data and build a unified picture, i.e. a mosaic of orthogonal yet complementary information. Therefore, efficient data analysis approaches incorporating the multivariate nature of various heterogeneous data sources is becoming more essential than ever.

To date, there are many computational approaches for multi-omics data integration (Huang et al, 2017). The existing solutions offer sample clustering and identification of genes and modules of genes with major contribution to the clustering results in the context of unsupervised analysis. Various supervised analysis methods are able to detect differentially expressed genes and form networks of molecular interactions consisting of those genes. Regardless of the nature of applications and the objectives of a given tool, there is little consensus on how data from multiple heterogeneous -omics platforms, observed at varying molecular levels, should be incorporated into the data analysis. While some methods treat individual molecular measurements as independent features of the data and explore the orthogonal feature subspace (Lin et al, 2013; Lock et al, 2013; Mo et al, 2013; Yuan et al, 2011), others attempt to model data transformation through a graph or kernel matrix using underlying intermolecular relationships (Bonnet et al, 2015; Vaske et al, 2010).

With an increasing number of *bona fide* genome-scale biological networks reported in the literature, network-based approaches offer another promising data integration strategy. Network-based methods identify modules of genes that are associated with a phenotypic outcome. They benefit from a synergetic combination of experimental data at hand and *a priori* information of functional relationships represented in the network data. Instead of relying on the numerical techniques to identify latent structures in the feature space, this line of approaches explicitly utilizes experimentally tested interactions as the background feature space in the analysis. Examples of network-based multi-omics data analysis include the latent-variable model of PARADIGM (Vaske et al, 2010), an unsupervised method to infer patient-specific pathway activation and deactivation status, jActiveModule (Ideker et al, 2002), a Cytoscape plugin that searches within a molecular network for expression activated subnetworks, and ATHENA (Kim et al, 2013), a tool for identifying complex susceptibility prediction models.

However, there is a significant gap in the ability of existing methods to connect two or more genes at different molecular levels. For example, some methods attempt to identify subnetworks of differentially expressed genes in each -omics platform first and integrate across different layers in a separate step (Bersanelli et al, 2016; Kim & Tagkopoulos, 2018). With the availability of more complete multi-omics data, we can now directly probe the relationship between different genes at multiple molecular levels. For instance, when we aim to test whether a gene encoding a transcription factor (TF) regulates the mRNA expression of a target gene, the most relevant data sources are the protein abundance for the TF gene and the mRNA abundances of its targets. When the goal is to test whether the interaction between protein molecules of two genes is elevated, the relevant omics data are the co-expression of the protein abundances of the two genes, not other -omics data such as mRNA abundances of those genes. To the best of our knowledge, no computational method connects multi-omics data at this level of resolution in the context of prediction analysis for phenotypic outcomes.

In this article, we present a new computational method iOmicsPASS, designed to integrate multi-omics profiles for phenotype prediction. We use biological networks as a workhorse to integrate data across different genes from different molecular levels and extract biological signals for the prediction of phenotypic outcomes. We perform two main tasks: (i) to integrate multi-omics data consisting of DNA copy number (optional), transcriptomics, and proteomics data by computing a consolidated table of co-expression scores, and (ii) to discover *predictive subnetworks*, a set of molecular interactions with co-expression patterns that best predict phenotypic outcomes.

The conceptual novelty of iOmicsPASS is clear: the method utilizes high confidence molecular interaction data as the basis of data linkage between different -omics platforms. Network data provides *a priori* knowledge for finding interactive features for predictive analysis. Moreover, the biological signals in the *predictive* subnetworks typically carry large effect sizes than gene signatures associated with the phenotype based on significance testing (*p*-value). The method is immediately generalizable to the integration of other -omics data using biological networks relevant to the given molecular types. Therefore, the analysis framework implemented in iOmicsPASS opens the door for other types of predictive network-based integration approaches based on co-expression of biomolecules.

## Results

### Overview of iOmicsPASS

The current implementation of iOmicsPASS takes quantitative transcriptomics and proteomics data and the biological networks linking the molecules as input. If DNA copy number is provided, then it is used to normalize the mRNA abundances. From these input data, we compute a co-expression score matrix for molecular interactions present in the network data. This *new co-expression data* is used in the subsequent analysis, where network features are selected for phenotype prediction.

In this work, we considered all biological networks as undirected graphs, where two interacting molecules (nodes) are connected with one another based on experimental evidence or high confidence computational predictions. These connections (edges) may include a pair of physically binding proteins or transcription/translation regulator (protein) and its target (mRNA). Our current implementation focuses on the integration of mRNA and protein data over a transcription factor (TF) regulatory network with or without normalization of mRNA abundance by DNA copy number data, and the integration of proteins over protein-protein interaction (PPI) network.

**Figure 1** summarizes the data analysis steps in iOmicsPASS. Overall, the iOmicsPASS workflow consists of four phases. In the first phase, we perform quality control and standardization of the individual -omics datasets to obtain Z-scores. In the second phase, we compute the edge-level co-expression data for biological interactions from related -omics data sets (node-level data). In the third phase (Subnetwork Discovery Module), the method selects predictive subnetworks using the modified nearest shrunken centroid (NSC) algorithm, originally implemented as the Prediction Analysis of Microarrays (PAM) (Tibshirani et al, 2002). Since PAM considers each feature (mRNA expression of one gene) as an independent variable, we modified the algorithm so that locally related molecular interactions tend to be selected together as predictive features. In other words, we modified the objective of the algorithm from asking “which genes are differentially regulated in group A?” to “which pairs of interacting genes are jointly differentially regulated in group A, considering other related genes?” In the last phase (Pathway Enrichment Module), we compute the enrichment scores for biological functions and pathways in the selected subnetwork for each phenotypic group (**Figure 1A**).

**Figure 1.**
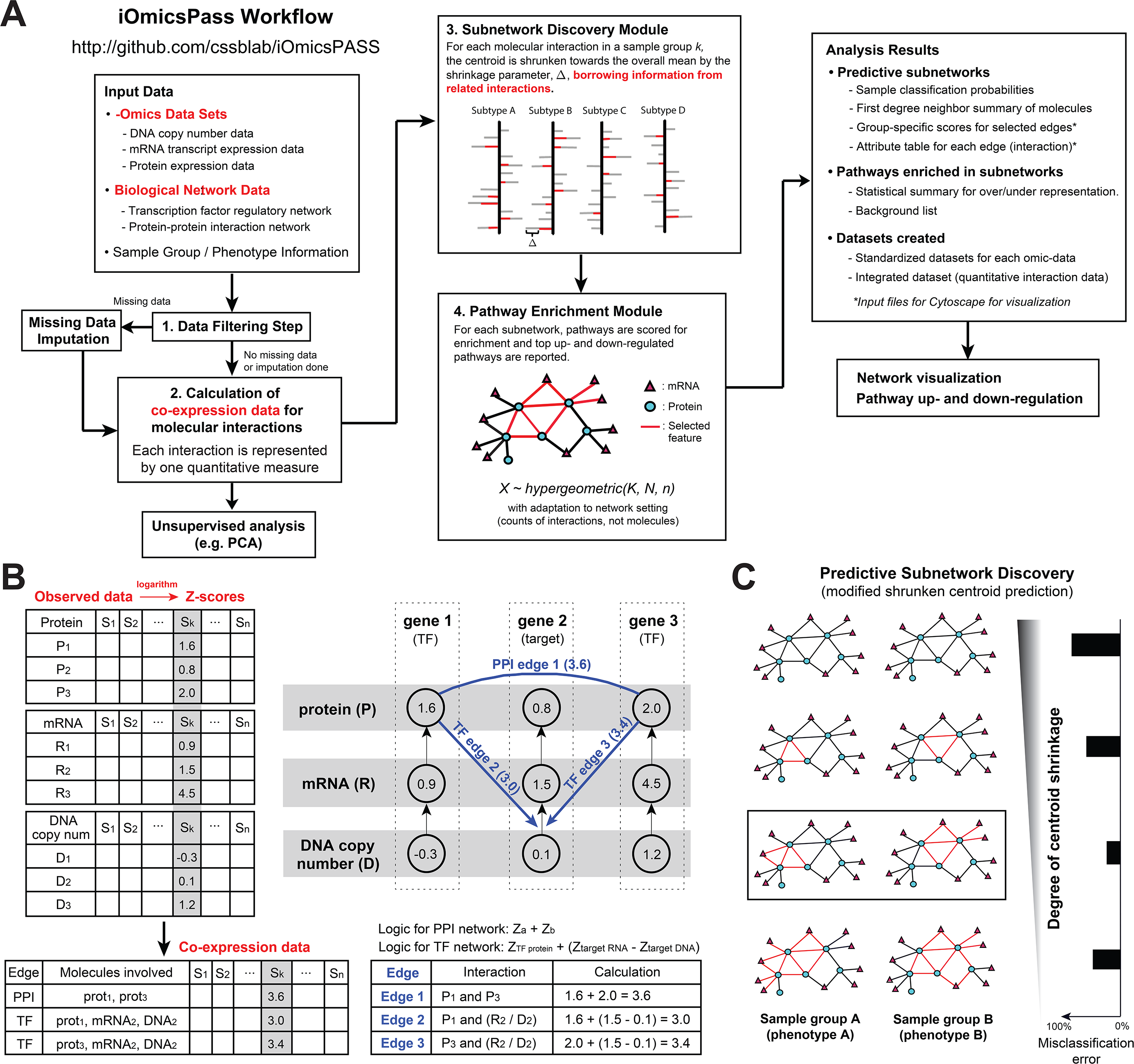
**(A)** iOmicsPASS workflow. iOmicsPASS takes multi-omics data, biological network data, and sample meta information. The molecular datasets go through quality control procedures and are integrated into an edge-level co-expression dataset for pairs of molecules as interactors in the biological interaction network. Predictive analysis module discovers the subnetwork signatures distinguishing phenotypic groups and associated biological processes. The software also produces a set of text files containing the details of the selected subnetworks and the materials for visualization of networks in the Cytoscape software. **(B)** The original multi-omics data is first standardized into Z-scores and converted to edge-level co-expression data. Shown is an example of two transcription factors (gene 1 and gene 3) and their common target gene (gene 2). Co-expression values are computed for the PPI between protein 1 and protein 3 and the transcription factor regulation between the two TF proteins and mRNA molecule of their target gene. **(C)** The resulting co-expression data is used as input to select the predictive edges for phenotypic groups using the modified nearest shrunken centroid algorithm.

The key first step in our workflow is to transform node-level data, i.e. original molecular abundance data, into edge-level data, i.e. co-expression scores of molecular interactions. Specifically, we derive a *co-expression* score for each pair of interacting genes (**Figure 1B**) based on the following logic. For an interaction between two genes (at the same or different molecular levels), a high co-expression score indicates that they have high chance of engaging in the biological activity, and vice versa. For example, if two physically interacting protein molecules have high Z-scores together, the sum of the two Z-scores will also be high, which indicates that there is increased chance of physical binding events.

Similar reasoning applies to two genes at different molecular levels. For example, if the protein abundance of a TF and the mRNA abundance of its target gene are both elevated in a phenotypic group, then the sum of Z-scores from both molecules will be high, indicating that the TF might regulate the respective target gene in that phenotypic group. In this case, the edge in the network describes a regulatory interaction, quantified by co-expression as above.

When DNA copy number is available, we subtract the Z-score of the copy number from the Z-score of the mRNA data of the same gene to normalize the abundance of the latter (per copy mRNA synthesis). Otherwise the high level of Z-score for mRNA may be entirely due to the copy number variation, not TF activity.

Next, we use the edge-level co-expression data as the input to the subnetwork discovery module. The module selects a subnetwork whose co-expression profile best predicts the membership of samples to each phenotypic group. To this end, we adopted the powerful NSC method called Prediction Analysis of Microarray (PAM) (**Figure 1C**), originally developed for gene expression microarray data (Tibshirani et al, 2002). Here we modified the NSC method so that the edges sharing common nodes tend to be selected together in the feature selection step, preserving the modular nature in the selected subnetwork. The subnetworks are selected using soft thresholding to minimize the misclassification error rates using cross-validations (see **Supplementary Information** for the detailed algorithm).

The analysis pipeline in iOmicsPASS produces several key results in separate tab delimited files: (i) a text-file containing edge-level co-expression data, (ii) a supervised classification model and shrunken centroid profiles for sample groups, (iii) data files to visualize the predictive subnetworks in Cytoscape, and (iv) a table of pathways enriched in each predictive subnetwork.

In summary, the novelty of iOmicsPASS is the intuitive transformation of multi-omics data into the co-expression scores of experimentally verified biological networks, a main backbone to connect interacting molecules within and across different molecular types. We also show that our integration approach is able to extract predictive information from each and every –omics platform in a balanced way. This is because the network information guides the method to combine horizontal (across different genes within the same molecular type) and vertical (across different molecular types and genes) information, instead of letting each data source compete with one another in the feature selection step.

### Simulation study: edge-level analysis outperforms node-level analysis in prediction accuracy

Network-level data analysis is thought to improve the sensitivity of identifying true signals (Wei & Li, 2007). However, previously reported evaluations have been made for the analysis of a single -omics platform, mostly mRNA gene expression data. The same hypothesis has not been thoroughly tested in the context of integrating multi-omics data. To objectively investigate the benefit of the data integration approach described above, we first performed simulation studies. Here we assumed the integration of transcriptomic and proteomic data using a TF regulatory network.

A key consideration in the simulation was the variable sensitivity of assay platforms. For example, the variations in the experimental parameters such as varying the number of PCR cycles for DNA quantifications introduce variable sequencing quality and distortion of quantification of molecules that results in loss of signal and detection of important molecules (Li & Stoneking, 2012; Meldrum et al, 2011; Meyer et al, 2008; Robin et al, 2016). Hence to account for the possibility of loss of signal during the assay quantification step, we planted signals in the proteins with a certain probability which we termed *assay sensitivity* (denoted as *P_AS_*) to reflect how likely the signal is observed in the experimental mass spectrometry (MS) data (**Figure 2A**). The question of interest is whether those proteins with undetected signals are recovered by the integrative analysis, which would otherwise be lost if the data were analyzed separately.

**Figure 2.**
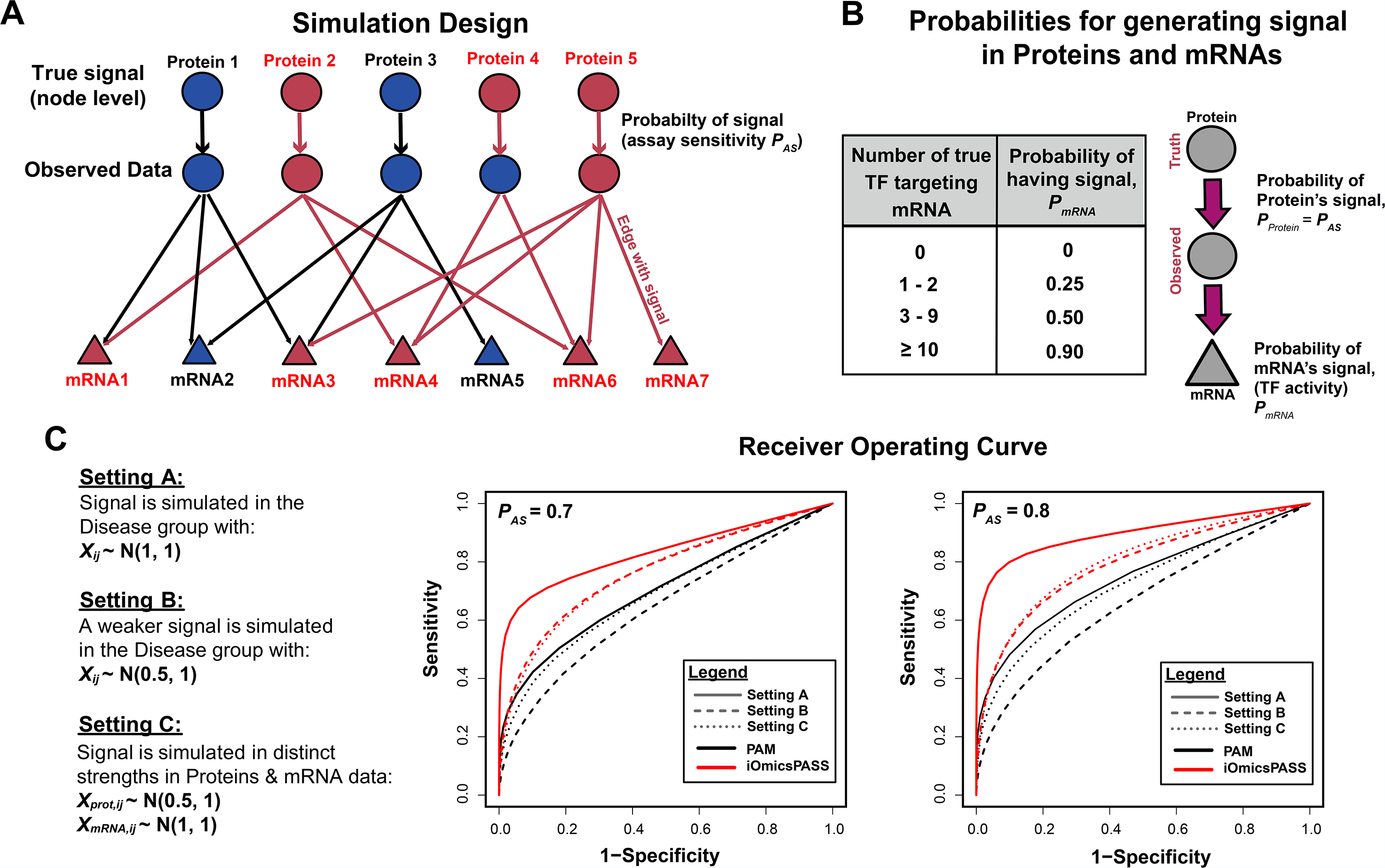
Simulation settings and results. (**A**) To reflect the noise level of MS-based proteomics, we simulate the protein-level data with assay sensitivity (P_AS_), i.e. the chance that the quantitative proteomic data shows group-to-group difference. (**B**) We also set the probability of increased transcription of mRNA molecules based on the number of active (more abundant) TF proteins targeting the gene (P_mRNA_). (**C**) Three different signal-to-noise ratio settings were used in the simulation (solid, dashed, and dotted lines). Red and black colors denote the performance of iOmicsPASS analysis on the derived co-expression data and that of PAM analysis on the matrix data formed by concatenating the two -omics data, respectively.

Another consideration in the simulation was that the homeostasis of the proteome is generally more tightly controlled than that of the transcriptome in dynamic cellular environment (Laurent et al, 2010). Moreover, untargeted MS-based, bottom-up proteomics still remains susceptible to a considerable amount of noise in the whole proteome profiling of complex tissues. Therefore, to reflect the reality that typical proteomic data shows more attenuated fold changes than transcriptomic data between different phenotypic groups, we applied different strengths of signals to the mRNA data and the protein data in one of the simulation setups (Setup C).

In each simulation setup, we compared the predictive analysis using the co-expression data as implemented in iOmicsPASS with a counterpart approach using the original PAM algorithm on the node-level data, assembled by concatenating individual -omics datasets into a single data matrix. Specifically, for the latter, we put mRNA and protein data into a single matrix and treated each molecule as an independent predictor of phenotype.

We generated a total of 100 sets of protein and mRNA data, each containing 1000 TF proteins and 5000 mRNAs of their target genes across 50 diseased and 50 control samples. We planted the signal, i.e. difference between the two phenotypic groups, in 100 randomly selected TF proteins and mRNAs of their target genes with stipulated probabilities in the samples of the disease group using Gaussian normal distribution with mean *μ*, and standard deviation *σ*. By contrast, the data for the samples in the non-disease group were generated from the background Gaussian distribution with mean 0 and standard deviation *σ*. **Figures 2A and 2B** illustrate the overall simulation design and probabilities used for generating signal in each molecule. Simulations were carried out using three different setups, varying the signal-to-noise ratios and the results were then averaged across the 100 sets in each setup (see **Methods** for details).

To construct the Receiver Operating Characteristics (ROC) curve, we calculated sensitivity and specificity in terms of the number of interactions, where we counted true and false by the number of edges, instead of nodes. We define true signal as an edge that connects any of the 100 selected proteins with its target mRNA molecules. Using the set of 1000 proteins and 5000 mRNAs that were simulated, a total of 48,682 possible edges can be constructed based on our curated network. Of those, we considered 6,742 (13.8%) edges as true signals. For iOmicsPASS, we considered the set of selected edges as predictive features. For PAM, as it selects each molecule as a predictive feature, we first formed edges between the selected nodes by the PAM algorithm (from the same network background) and counted the edges as predictive features by PAM.

**Figure 2C** shows the comparison between the node-level analysis using the PAM algorithm and the edge-level analysis using iOmicsPASS, when the signal-to-noise ratio and probability of *assay sensitivity* (*P_AS_*) are varied. The ROC curves clearly show that the edge-level prediction analysis outperforms the node-level analysis in terms of both sensitivity and specificity in all three simulations setups. At *P_AS_*=0.7, the area under the curves (AUC) for all three setups using iOmicsPASS were 10.8% to 19.3% higher than that compared to the results using PAM (**Table 1**). The difference in AUC is the largest in setting A (PAM: AUC = 0.701; iOmicsPASS: AUC = 0.832), where the planted signal was the strongest among all setups (i.e. *μ*=1). The performance is similar in both settings B and C when using iOmicsPASS with AUC values of 0.765 and 0.761, respectively. But the node-level analysis using PAM consistently performed the worst in setting B with the smallest AUC (AUC =0.641) when the signal is noisier than in setting A (i.e. *μ*=0.5). Overall, our simulation studies suggest that there is clear benefit in integrating data as co-expression data over networks as proposed in iOmicsPASS.

**Table 1.** The area under the curve values for the three different simulation setups comparing node-level analysis using PAM and edge-level analysis using iOmicsPASS.

| Area under the curve<br>(AUC) | <u>$P_{AS} = 0.7$</u> |  | <u>$P_{AS} = 0.8$</u> |  |
| --- | --- | --- | --- | --- |
|  | PAM | iOmicsPASS | PAM | iOmicsPASS |
| **Simulation Setup A** | 0.701 | 0.832 | 0.744 | 0.897 |
| **Simulation Setup B** | 0.641 | 0.765 | 0.659 | 0.790 |
| **Simulation Setup C** | 0.687 | 0.761 | 0.721 | 0.801 |

Next, we tested the performance of iOmicsPASS in a more realistic scenario: when network information is incomplete and noisy, i.e. contains many spurious interactions (false positives) and lacks many potential *bona fide* interactions (false negatives). To this end, we further simulated *noisy networks* that include spurious interactions and lack a portion of true interactions (see **Supplementary Information**). The results show that, even without a complete network, the edge-level analysis using noisy networks still outperforms that of the node-level analysis in all three simulation setups. In the setup C, we still observed the smallest AUC of the ROC curves in the results by the node-level analysis even though the overall prediction accuracy was similar to each other. Therefore, we argue that there is significant value in adopting an edge-level prediction analysis approach, i.e. using network information, as long as the biological network data at hand captures the key molecular interactions in the true network.

### Analysis of breast cancer subtypes in TCGA BRCA cohort

Next, we assessed the ability of iOmicsPASS to discover predictive subnetworks for breast cancer subtypes. We used the invasive ductal breast carcinoma (BRCA) data of TCGA as a benchmark cancer multi-omics data, which contains the well-defined four intrinsic subtypes for breast cancer based on the mRNA-based PAM50 signature (Parker et al, 2009). In general, the subtypes are characterized by the immunohistochemistry of three receptor proteins: estrogen receptor (ESR1), progesterone receptor (PGR), and human epidermal growth factor 2 (HER2/neu, also known as Erb-b2). The two luminal subtypes (i.e. Luminal A and Luminal B) have positive expression of ESR1 and PGR. HER2-enriched (HER2E) subtype has chromosomal amplification-associated over-expression of HER2 protein. The Basal-like subtype, often characterized by negative expression of ER, PR, and HER2, is associated with the poorest prognostic outcome, and patients in this subtype largely remain without clear treatment options beyond chemotherapy. We used the PAM50 classification of the samples as true class labels in the prediction analysis. Our goal here was to evaluate the ability of iOmicsPASS to correctly classify the tumors to the predefined subtypes, while identifying a TF regulatory network and a PPI network uniquely representative of each subtype.

#### Edge-level co-expression data analysis has advantages in the integration of heterogeneous molecular data

We integrated proteomics, transcriptomics, and DNA copy number data, produced by the CPTAC (Mertins et al, 2016) and TCGA (Cancer Genome Atlas, 2012), respectively. TCGA’s original multi-omics data has a total of 1,098 tumor samples, but only 105 samples were analyzed in the CPTAC’s proteomics profiling. Among these, 103 samples had DNA copy number data, transcriptomics and proteomics data available. TCGA assigned 24 to Basal-like, 18 to HER2E, 29 were Luminal A and 32 were Luminal B subtype based on mRNA data, respectively. In our analysis, we used transcriptomics and proteomics data since this analysis provided the lower test classification errors than the one incorporating DNA copy number data.

The subnetwork discovery module identified subtype specific subnetworks consisting of 3,453 edges, using a threshold that keeps the misclassification error below 30%. **Supplementary Table 1** provides detailed information on the selected subnetwork. **Figure 3A** shows the principal component analysis (PCA) plot of the co-expression scores, with annotation of misclassified tumor samples. Although the overall classification accuracy is above 70%, about half of the HER2E tumors are grouped with Luminal A tumors and the other half with Luminal B tumors in this analysis, mostly along the PC1 axis. **Figure 3B** also shows that the prediction had produced the highest test errors for the HER2E subtype. The “training” errors for HER2E and Luminal B were also the highest in our analysis (55.6% and 34.4% respectively).

**Figure 3.**
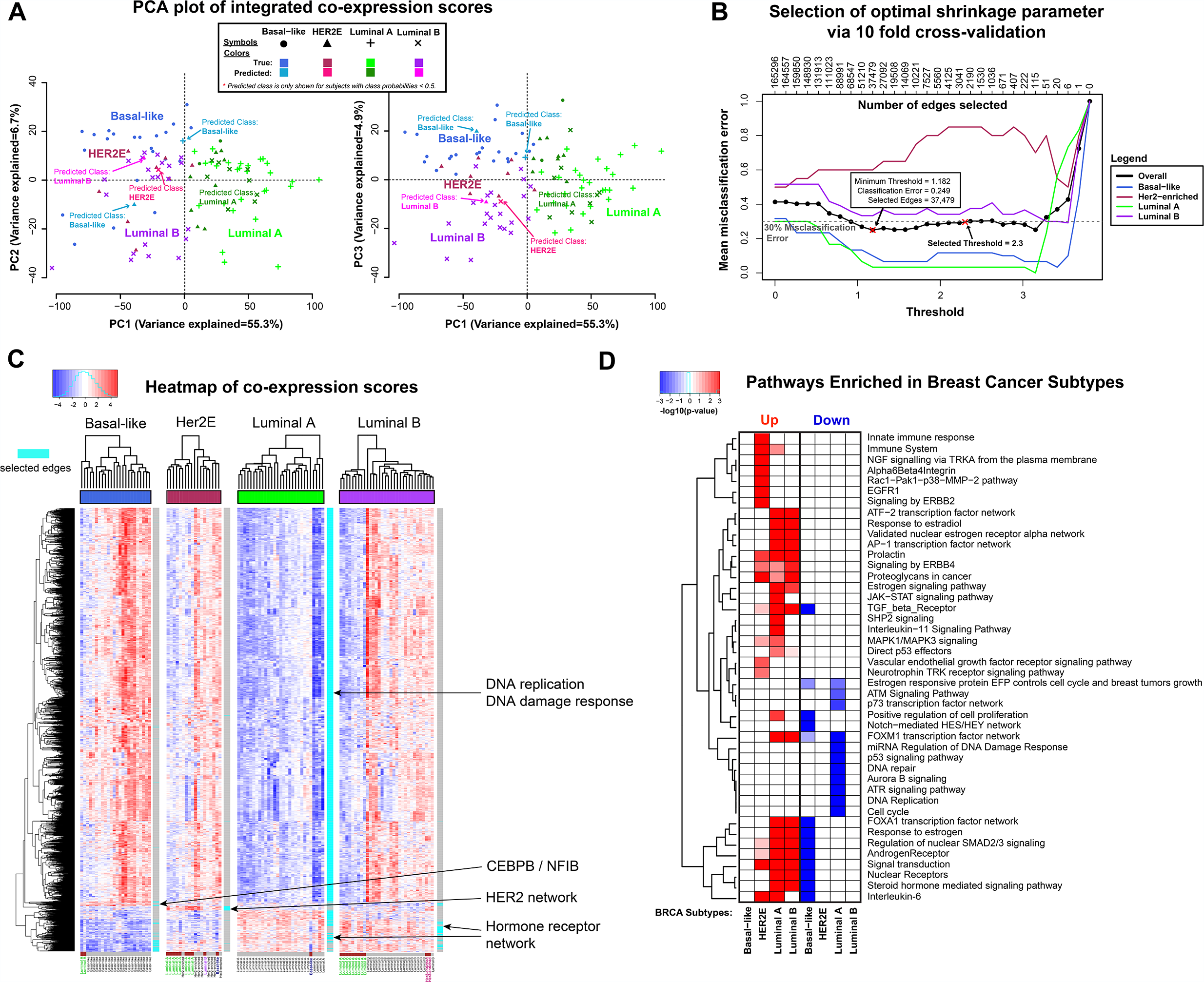
BRCA data analysis. (**A**) PCA plots for (PC1, PC2) and (PC1, PC3) pairs. Tumor samples misclassified by the optimal prediction model, trained on the same data, are highlighted. Each symbol is the known subtype from mRNA-based PAM50 classification (Parker et al, 2009) provided in TCGA. The symbols were colored in two different shades, depending on whether predicted class matches the known subtype or not. (**B**) Misclassification errors (test errors) computed for each subtype using 10-fold cross-validation. The red color cross on the left side is the threshold that gives the smallest misclassification error rate, and the one on the right side is the threshold giving a sparser subnetwork within one standard deviation rule in the curve (rule of thumb). (**C**) Heatmap of the co-expression data for the union of all four subnetworks. The cyan color bar on the right-side of the heatmap highlights the portion of the subtype-specific subnetworks. (**D**) Heatmap of statistical significance for the enrichment of pathways in the subtype specific co-expression signatures. Shown in the diagram are minus the logarithm of p-value (base 10).

**Figure 3C**, the heatmap of co-expression data, shows why the integrated data does not completely align with the PAM50-designated subtypes. Although HER2E tumors demonstrate the well-known up-regulation of ERBB2 and GRB7 protein expression, the more dominant profile in the overall predictive subnetwork is that of the DNA replication and DNA damage response network, dividing the HER2E tumors into two sub-clusters resembling Luminal A and Luminal B subtypes. The split pattern of DNA replication and DNA damage response network is also visible for Luminal B tumors. About two quarters of those samples have clear elevation of co-expression of DNA replication genes, while the remaining samples show low co-expression, leading to misclassification of those tumors into Luminal A subtype.

This observation led us to hypothesize that the inclusion of DNA replication and damage response markers such as the members of mini-chromosome maintenance (MCM) complex, in addition to the hormone receptor status, is an important axis of information in breast cancer subtyping in all subtypes except Luminal A. This interpretation is also supported by recent reports that alternative genes such as MCM2 can be better prognostic markers than the current markers of cell proliferation such as the percentage of tumor cells expressing Ki-67, a key criterion used for deciding between Luminal B and Luminal A subtypes (Yousef et al, 2017).

Next, we compared the classification performance between the edge-level analysis and the node-level analysis, where we concatenated the three -omics data sets into a single matrix and applied the powerful PAM method. We recall that the molecular subtypes were designated using the mRNA data, and thus the node-level analysis is expected to slightly outperform the edge-level analysis. Indeed, **Supplementary Figure 2B** shows that the misclassification rates for Luminal B and HER2E subtypes were smaller for the node-level analysis. However, when we looked at the predictive features in the node-level analysis, more than 95% of the predictive features were mRNA molecules. Despite seemingly better prediction performance, the concatenation-based integration almost completely fails to incorporate additional predictive information provided by the proteomic data – the very molecular level where the expression of functionally active gene products is observed. By contrast, our analysis framework utilizes the biological network as the backbone of the feature selection, which guides the predictive subnetwork selection to incorporate both molecular types in a more balanced manner.

#### Predictive analysis reports three network axes dividing the subtypes

The predictive subnetworks reported by iOmicsPASS are stored in “EdgesSelected_minThres.txt” file, and one can directly import this file into the Cytoscape tool to produce a network diagram. See **Figure 4** for Cytoscape visualization of the integrated - omics data for the four subnetworks, shown in a common background network (union of the four).

**Figure 4.**
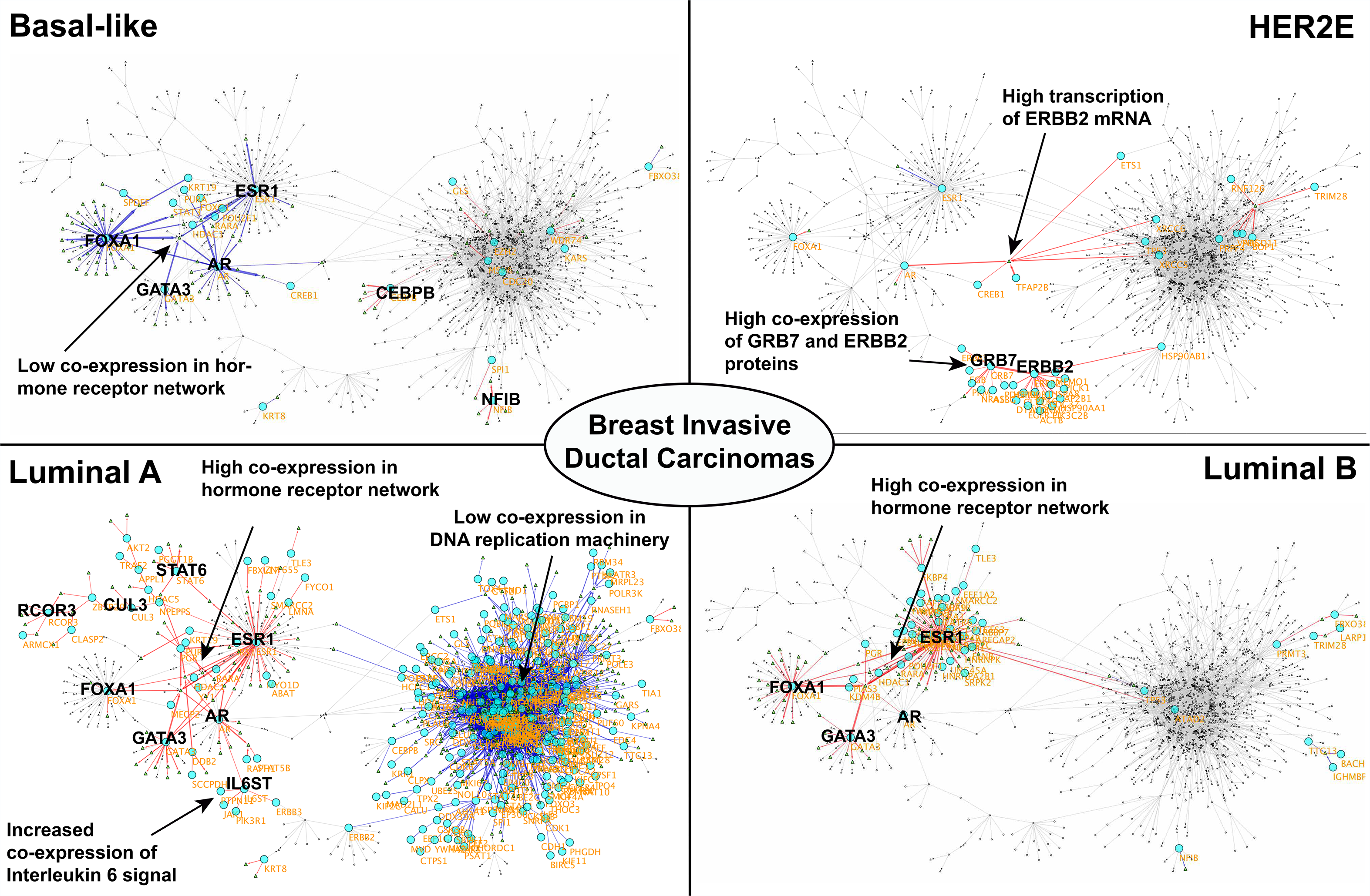
Subnetwork signatures of four subtypes in BRCA illustrated in Cytoscape. Red and blue lines (edges) are molecular interactions (TF regulation or PPI) with higher and lower co-expression values compared to the overall centroid, i.e. average profile in the current data set. Proteins and mRNA molecules are shown as large circles and small triangles, respectively. The thickness of an edge is proportional to the absolute co-expression value.

The subnetwork discovery module revealed that there are *three major axes* segregating the samples into the four molecular subtypes. First, a large subnetwork consisting of the proteins involved in DNA replication and DNA damage response is down-regulated (relative to other tumors) in terms of TF regulation and PPIs in Luminal A subtype (blue edges), relative to the average breast tumors in the study. This finding is consistent with our expectation since Luminal A is the group of breast cancer patients with the best prognostic outcome (Parker et al, 2009).

The second axis is a subnetwork of hormone receptors and related proteins, consisting of protein molecules such as estrogen receptor (ESR1), androgen receptor (AR), and forkhead box protein A1 (FOXA1) genes, which are particularly up-regulated in the both Luminal subtypes and down-regulated in the Basal subtype according to the immunohistochemistry data from TCGA. The third axis consists of physical interactome of protein molecules of HER2 and growth factor receptor bound protein 7 (GRB7) for the HER2E subtype. GBR7 is a SH2-domain adaptor protein binding to receptor tyrosine kinases, and it is itself a proto-oncogene co-amplified with ERBB2 (HER2/Neu) in breast cancer (Pandey et al, 1996; Vinatzer et al, 2005).

Interestingly, iOmicsPASS identified CCAAT/enhancer-binding protein beta (CEBPB) and nuclear factor I B-type (NFIB) as over-expressed protein molecules with direct impact on mRNA transcripts in the Basal subtype. Basal subtypes have long been characterized by negatively expressed markers (e.g. ER, PR, HER negative in triple negative cases), and characterization of CEBPB protein isoform in aggressive carcinomas in breast and other cancers have shown emerging literature evidence (Zahnow, 2009). Over-expression of NFIB has also been linked to poor prognostic outcomes in ER-negative breast tumors in the literature (Becker-Santos et al, 2017; Takeno & Sakai, 1991). Therefore, the findings warrant further investigation for the discovery of positively expressed markers for the Basal subtype.

From these three axes, we can also deduce that deterministically assigning each patient to a single subtype is not as straightforward as it may seem from the mRNA-based classification (PAM50), consistent with a previous working group report of oncologists (Guiu et al, 2012). Instead, our analysis suggests that subtyping should be done in a combinatorial manner: one person’s tumor may share molecular characteristics of two or more subtypes. While both Luminal A and B subtypes are indeed mostly ER positive, the major difference between the two is the reduced co-expression of DNA replication machinery in the Luminal A subtype. This data also explains why tumor samples labeled as HER2 subtype by PAM50 are not completely separable from those labeled as Luminal B subtypes (**Figure 3C**). In fact, some subjects classified under Luminal B subtype exhibited positive HER2 expression as well. The dichotomous subgrouping is clearly visible in the co-expression profiles, where each subtype shows heterogeneity in the subnetwork representing DNA replication machinery and damage response.

#### Pathways underlying subtype specific subnetworks

Lastly, iOmicsPASS tests for the enrichment of biological functions and pathways in the predictive subnetworks, using hypergeometric test on the edge counts (see **Supplementary Information** for the detailed description). **Figure 3D** shows the summary of pathway enrichment in the subtype-specific subnetworks. As expected, the two Luminal subtypes share similar functional enrichment profiles in the up-regulated and down-regulated edges, including hormone receptor and signal transduction pathways as well as FOXA1 and AP1 transcription factor network. HER2E subtype shows over-representation of cell signaling pathways not up-regulated in the Luminal subtypes, including epidermal growth factor receptor (EGFR) and platelet-derived growth factor (PDGF), and elevated immune response. Luminal A subtype was largely characterized by the entire cell proliferation machinery down-regulated relative to the other subtypes. At the pathway level, the Basal subtype did not show any up-regulated pathways as expected from the past literature.

A caveat, however, is that the data were integrated in the space of TF network and PPI network, which currently ignores other -omics data available for these samples, including DNA methylation, microRNA, and phospho-proteomic data (Mertins et al, 2016). Incorporation of those data, through calculation of meaningful co-expression measures for related networks (e.g., connecting them through the respective genes), is likely to segregate samples into finer groups and potentially identify additional key subnetworks in the future.

### Analysis of colorectal cancer subtypes in TCGA CRC cohort

We next applied iOmicsPASS to the multi-omics data for subjects in the colon and rectum adenocarcinoma cohort (CRC) from TCGA and CPTAC. Unlike breast cancer, the molecular subtypes in CRC are less characterized with respect to therapeutic targets (e.g. ER, HER2 in breast cancer), and thus integrated molecular subnetworks may provide more insight to subtype-specific therapeutic target identification.

Similar to BRCA analysis, we aimed to identify predictive subnetworks for three subtypes TCGA assigned based on the transcriptomic signature from De Sousa e Melo *et al* (De Sousa et al, 2013). This subtyping method divides tumors into three groups: chromosomal instable subtype (CIN), microsatellite instability/CpG island methylator phenotype (MSI/CIMP), and a third subtype called Invasive subtype, characterized by up-regulation of genes involved in extracellular matrix remodeling and epithelial-mesenchymal transition (EMT), a signature of sessile-serrated adenomas.

A total of 633 tumor samples (276 individuals) from colon and rectal cancer patients were profiled by TCGA, and among those the proteomes of 95 samples were profiled by the CPTAC. Out of 220 individuals with expression subtyping information assigned by TCGA, 88 belonged to CIN, 71 to MSI/CIMP, and 61 to Invasive subtypes, respectively. In this analysis, we incorporated DNA copy number as the normalizing constant to mRNA data as illustrated in **Figure 1B**, since this gave better predictive performance in terms of misclassification errors (comparative results not shown).

**Figure 5A** shows the PCA analysis of the edge-level co-expression data for the selected subnetworks, which consists of 1,991 biological interactions. See **Supplementary Table 2** for the detailed information on the subnetwork. As we observed from the BRCA data, the first two PCs clearly show the systemic similarity and dissimilarity between the three subtypes. Consistent with the characterization in De Sousa e Melo *et al*. (De Sousa et al, 2013), the MSI/CIMP subtype is distinguished from the other two subtypes along PC1, which explains 24.6% of the total variability among the samples. PC2 divides the CIN and Invasive subtype, although some samples designated as the CIN subtype were grouped together with the Invasive subtype.

**Figure 5.**
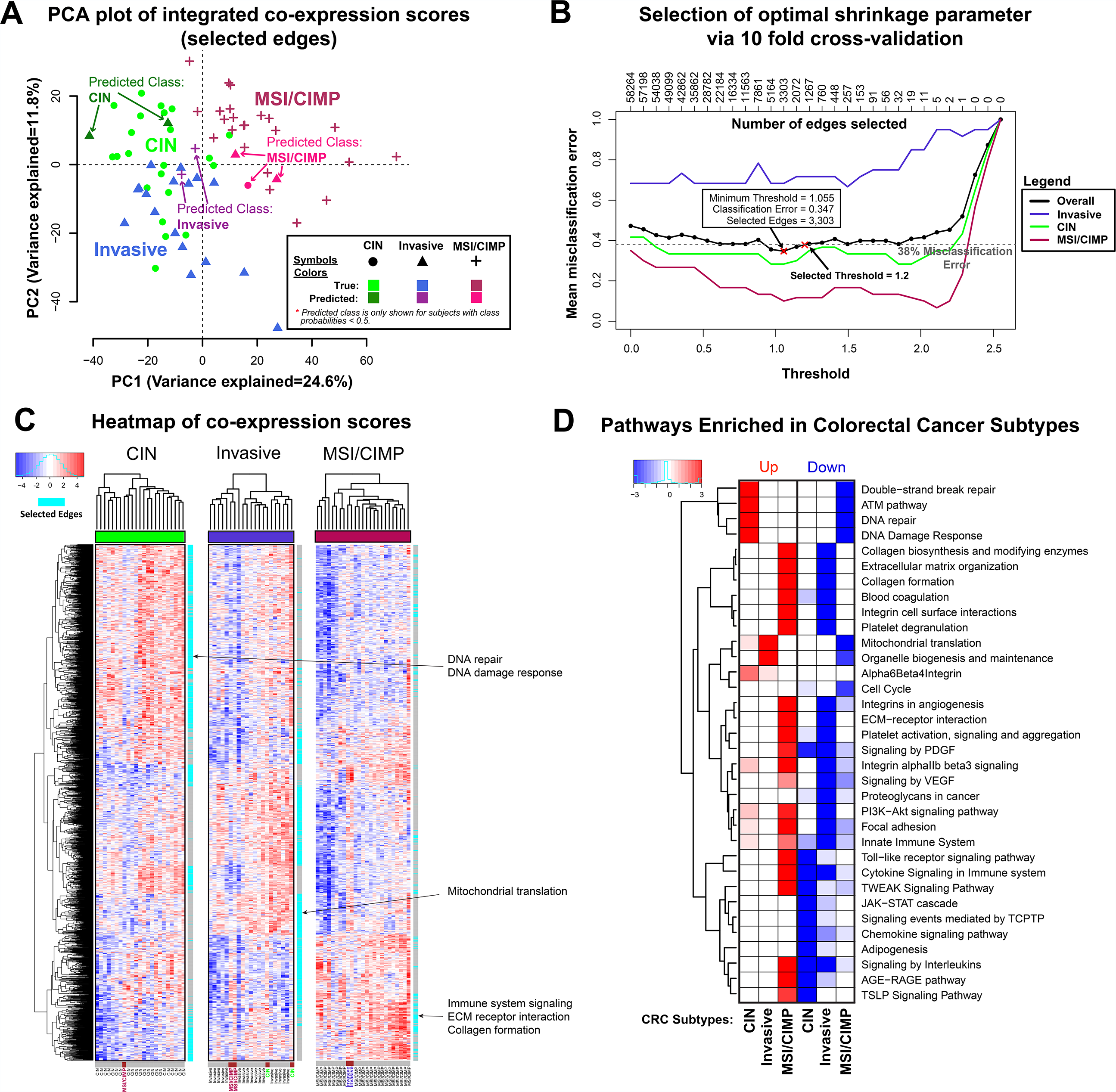
CRC data analysis. (**A**) PCA plots for PC1 and PC2. Tumor samples misclassified by the optimal prediction model, trained on the same data, are highlighted. Each symbol is the known subtype from mRNA-based classification (De Sousa et al, 2013) provided in TCGA. The symbols were colored in two different shades, depending on whether predicted class matches the known subtype or not. (**B**) Misclassification errors (test errors) computed for each subtype using 10-fold cross-validation. The red-colour cross on the left side is the threshold that gives the smallest misclassification error rate, and the one on the right side is the threshold giving a sparser subnetwork within one standard deviation rule in the curve (rule of thumb). (**C**) Heatmap of the co-expression data for the union of all four subnetworks. The cyan-colour bar on the right-side of the heatmap highlights the portion of the subtype-specific subnetworks. (**D**) Heatmap of statistical significance for the enrichment of pathways in the subtype specific co-expression signatures. Shown in the diagram are minus the logarithm of p-value (base 10).

#### Edge-level analysis gives a more balanced signature of transcriptome-proteome than node-level analysis

Next, the subnetwork discovery module reported a total of 1,991 predictive edges across the three subtypes. The estimated misclassification error rate, i.e. the test error based on cross-validation (**Figure 5B**), was the smallest for the MSI/CIMP subtype, as the transcriptional co-expression signature of signal transducer and activator of transcription protein family proteins (STATs) and their target transcripts clearly separated them from the other two groups (**Figure 5C**). The Invasive subtype was the class with the highest test error, as a large number of those tumors actually had high co-expression of DNA repair and DNA damage response molecules and low co-expression of immune response-related genes typically observed in the CIN subtype, consistent with the PCA plot (**Figure 5A**).

We again compared the classification performance and the composition of predictive features between the edge-level analysis and the node-level analysis. **Supplementary Figure 3** shows the test error plot using PAM based on a ten-fold cross-validation. Similar to the experience in the BRCA data, the overall misclassification error rates tend to be slightly lower in the node-level analysis than iOmicsPASS. However, the predictive features were again dominated by the mRNAs (77.9%), and many of these mRNA signals did not show consistent signals at the protein level when it was quantified. When the proteomic data is incorporated through iOmicsPASS, the TF regulatory connections and physical interactome produced sample classification results inconsistent with the result solely based on the transcriptomic data.

#### Subnetwork discovery module reveals four axes of predictive subnetworks

The network diagram of integrated data in **Figure 6** suggests that the predictive subnetworks have several major axes in the network structure. The first axis is the large PPI and TF subnetworks involved in DNA damage, DNA repair, and mRNA splicing/spliceosome, which is the most up-regulated in the CIN subtype (middle of the left panel). The second axis is the PPI network of ribonucleoproteins, especially those implicated in protein translation and organelle biogenesis, which are up-regulated in the Invasive subtype (lower right of the middle panel). The Invasive subtype was also characterized by low co-expression for the PPI network of collagen subunits and other glycoproteins in the extracellular matrix (ECM) (upper left of the middle panel). The last axis is the pro-inflammatory, immune signaling subnetwork represented by signal transducer and activator of transcription protein family (STATs) and p65 (RELA), up-regulated in the MSI/CIMP subtype (lower left of the right panel).

**Figure 6.**
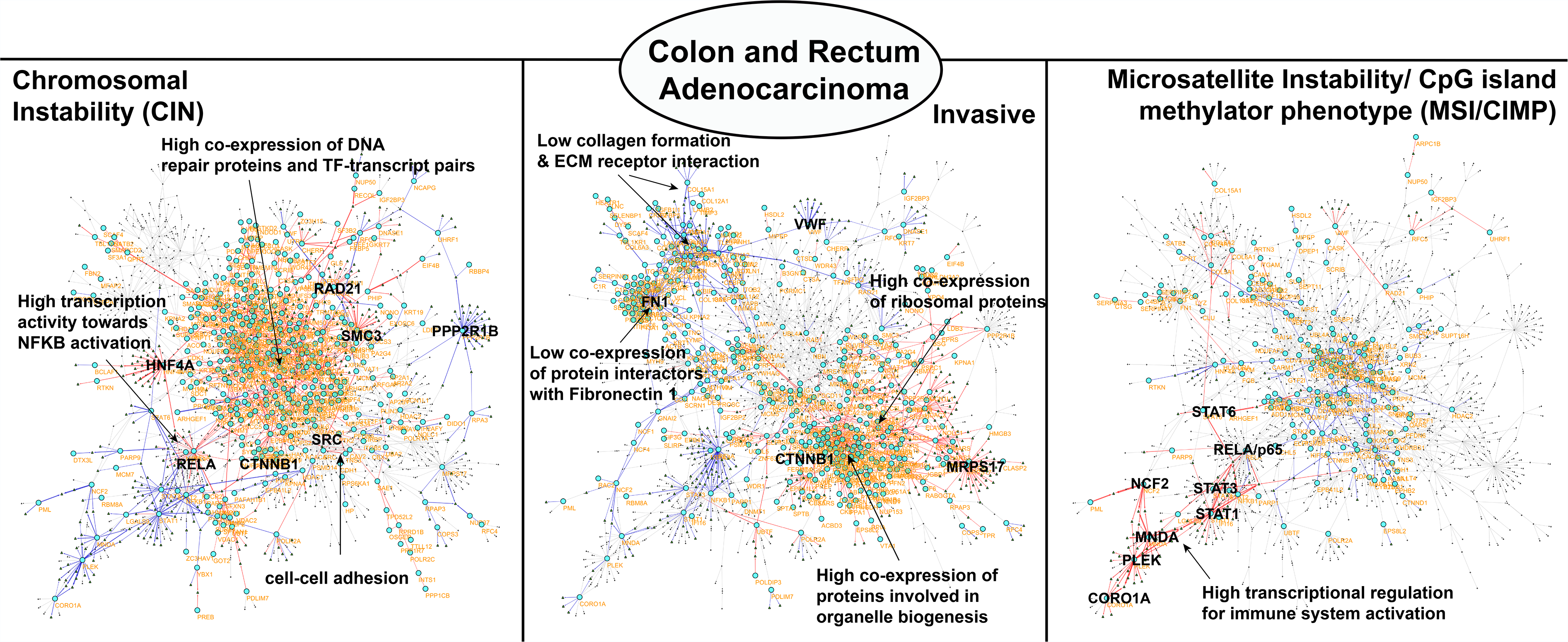
Subnetwork signatures of four subtypes in CRC illustrated in Cytoscape. Red and blue lines (edges) are molecular interactions (TF regulation or PPI) with higher and lower co-expression values compared to the overall centroid, i.e. average profile in the current data set. Proteins and mRNA molecules are shown as large circles and small triangles, respectively. The thickness of an edge is proportional to the absolute co-expression value.

The pathway enrichment analysis confirmed the enrichment of the aforementioned biological processes in the predictive subnetworks (**Figure 5D**). The subnetwork signature for the CIN subtype is markedly represented by up-regulation of DNA repair and DNA damage response and down-regulation of immune response; the MSI/CIMP subtype shows up-regulation of immune response through cytokine/chemokine/interleukin signaling and other signaling pathways; and lastly, the Invasive subtype is characterized by up-regulation of mitochondrial translation and organelle biogenesis and maintenance, possibly compensating for massive energy expenditure required for cell proliferation and mobility, and down-regulation of signaling pathways and cellular structural component regulation (e.g. ECM organization, collagen formation, platelet degranulation and activation).

#### Key oncogenic protein hubs in the CRC subtypes

We investigated the predictive subnetworks in greater details, with respect to the key network hub proteins (**Supplementary Table 2**). First, the MSI/CIMP subtype has elevated co-expression of proteins present in the extracellular space, including platelet aggregation, hemostasis (FN1, FGA, FGB, SERPINF2, PLG) and immune response (C1QB, C4BPA, CLU, FGA, FGB, S100A9, PML), i.e. pro-inflammatory markers may have to do with platelet aggregation near tumor lesions. Among the transcription factors, STAT1/2/3, MNDA, PLEK, NFKB1 and PARP1 were up-regulated with a majority of their target genes’ mRNAs consistently up-regulated in this subtype.

Next, we examined the predictive subnetwork for the CIN subtype. The large subnetwork in the CIN subtype, shown in the middle part of CIN subnetwork (**Figure 6**), consists of multiple components including pre-mRNA processing RNA-binding proteins (e.g. helicases) and splicing factors, and mitochondrial ribosomal proteins, and 14-3-3 protein complexes. As expected, this network also contained DNA repair and DNA damage markers such as RAD21/RAD50, structural maintenance gene SMC3, p53-binding protein TP53BP1, and mismatch repair gene MSH6. Another part of the protein interactome subnetwork, located in the lower right part of the network (lower right part in the CIN subnetwork, **Figure 6**), was that of cell-cell adhesion containing protein interactions between proto-oncogene SRC, catenin beta 1 CTNNB1, and cadherin CDH1. However, the co-expression in this network was also higher in the Invasive subtype than the MSI/CIMP subtype.

In the TF network, a number of protein molecules of TF genes were up-regulated in the CIN subtype. Hepatocyte nuclear factor 4 (HNF4) and translocase of outer mitochondrial membrane 34 (TOMM34), previously associated with chromosome 20q amplicon and elevated protein-level abundance (Zhang et al, 2014), are more pronounced in the CIN subtype over the others.

Lastly, the subnetwork with high co-expression in the Invasive subtype consisted of many ribosomal proteins, ribonucleoproteins, and mitochondrial translation genes, indicating that enhanced ribosome biogenesis is a key marker of this subtype in CRC (Lai & Xu, 2007). The current literature also suggests that the Invasive subtype is expected to have elements of WNT signaling pathways and epithelial-mesenchymal transition (EMT) (Vu & Datta, 2017). Unfortunately, the protein-level data of adenomatous polyposis coli (APC) gene, an important hub, did not qualify and was excluded from the prediction analysis. In addition, we observed elevated co-expression for the SRC and ephrin type B receptor 2 (EPHB2), a substrate of SRC that increases cell invasiveness via tyrosine phosphorylation (Leroy et al, 2009; Pasquale, 2010), which can be interrogated as a prime therapeutic target for the Invasive subtype.

#### Addressing the ambiguities in CRC subtypes

Our edge-level analysis of co-expression data also unveils the complexity underlying the current CRC subtyping. While MSI/CIMP subtype was easily distinguishable from the others, the integrated data showed mixed clustering results for the other two subtypes. As we observed in the case of HER2E and Luminal B subtypes in BRCA, the TF regulatory network and interactome for the Invasive subtype share those of CIN subtype tumors, suggesting that some tumor samples have molecular characteristics of both subtypes at the same time.

Although this observation again points to further investigation of shared features of predictive subnetworks between the two subtypes, we conclude that the data integration through biological networks was able to incorporate different -omic data in a balanced way, revealing an extended level of heterogeneity at the systemic level. This observation underscores the need for more refined subtyping and identification of therapeutic markers in the newly emerging clusters of CRC patients. Incidentally, a more recent report drew consensus molecular subtyping (CMS1-4) from multiple studies (Guinney et al, 2015) with further supporting evidence from in vitro and in vivo models (Linnekamp et al, 2018), yet this information was not incorporated into TCGA’s sample manifest information.

## Discussion

In this work, we presented a computational method for integrating multi-omics data in the space of biological networks and extracting a unique network signature predictive of each phenotypic group. The key difference of iOmicsPASS framework from other integration approaches is that the molecular data that are pertinent to the given network are integrated through the creation of co-expression scores. The co-expression scores effectively represent the potential for the interactions between template and product molecules, physically interacting partners, regulators and their targets. Our analysis framework represents a conceptual departure from the classical node-level analysis methods, where network-level knowledge is used as a *post hoc* assembly of significant findings. In addition, iOmicsPASS substantially reduces data dimensionality by extracting a *predictive* subnetwork from the entire network provided by the user, where the optimal subnetwork minimizes the overall sample misclassification errors. In our experience, this selection method requires stronger signals, i.e. greater magnitude of differences, than other methods reliant on statistical hypothesis testing (*p*-values for associations).

The proposed data analysis framework can be immediately generalized to the integration of other types of data, including integration of DNA methylation with mRNA / DNA copy number data to find epigenetic signatures that has impact on the transcriptional activities, microRNA with protein / mRNA data to detect miRNA-driven post-transcriptional activities in different sample groups. In both examples, one molecular type of molecules (e.g. miRNA) modulates the synthesis of the other type of molecules (e.g. protein per mRNA copy), the calculation of edge-level co-expression data must be adapted to the given relationship between molecules.

In the case of integrating mRNA data and DNA methylation in the regulatory regions or gene bodies of the coding genes, the relevant biological network can be constructed based on the proximity of methylation locus to the transcription start site of the nearest gene in the mRNA data. The modification suppresses the transcript output of the gene for the methylation of regulatory regions, and thus the co-expression by the sum of the two standardized Z-scores has to be changed to the difference between the former (mRNA) to the latter (DNA methylation). By contrast, the modification on gene bodies increases the output, and thus the co-expression should remain as the sum of the two Z-scores (Yang et al, 2014). The same principle also applies to the integration of miRNAs by the difference between the protein score and the mRNA score, where the output of protein per mRNA molecule is negatively modulated by the microRNA activity.

Meanwhile, the method has ample room for future extensions. First, the current implementation does not allow for prediction of phenotypic groups for external datasets. However, this was a deliberate choice: it is difficult to imagine typical molecular profiling studies with all three -omic platforms universally applied to hundreds of tumor samples, especially when proteomics is a part of the repertoire. Hence our immediate future work is to construct a framework to translate the predictive subnetworks into *a prediction machine* for a new dataset with *incomplete multi-omics data*.

Second, as we allow integration of more diverse type of data sets, some -omics data will wield more influence than others, as we saw in the examples of HER2E and Invasive subtypes. With further extensions to other -omics data types and additional modules for prediction, we believe iOmicsPASS offers one of the most comprehensive multi-omics data integration methods in the current literature.

Lastly, our analysis showed that the molecular subtypes are not mutually exclusive. The current practice of molecular subtyping is to let all subtypes to compete for assignment for a tumor sample. In other words, there is only one winner in the assignment. As we demonstrated in both cancer datasets, there will be a subset of tumor samples that share characteristics of multiple subtypes, given typical heterogeneity in tumors. To address such cases, future prediction method will have to embrace the multiplicity of assignments, which will eventually be a determining factor in the choice of therapeutics for specific tumors.

## Material and Methods

### Development of iOmicsPASS

iOmicsPASS is a bioinformatics workflow with four phases of data analysis. In the first phase, we remove molecules with many missing data points in each -omics dataset and use the weighted *K*-nearest neighbor method (Schwender, 2012) to impute the missing data before Z-score standardization.

In the second phase, the node-level quantitative data is transformed into an integrated set of co-expression measurements (edge-level) for every pair of interacting molecules in the biological network. For PPI networks, the sum of the standardized log-scale (base 2) measurements of two interacting proteins forms a co-expression measurement for each protein-protein interaction (edge). For TF regulatory networks, the sum of the standardized log-scale (base 2) Z-score of a transcription factor protein and Z-score of the mRNA of its gene target forms a co-expression of protein-mRNA edge, where the latter element can be further normalized by the Z-score of the DNA copy number variation (if available) to represent mRNA abundance per DNA copy.

In the third phase, we apply the nearest shrunken centroid (NSC) algorithm of the Prediction Analysis of Microarray (PAM) to the integrated co-expression dataset and report the predictive subnetworks. We call the collection of phenotypic group-specific subnetworks as a subnetwork signature throughout the manuscript. We also modified the NSC algorithm to account for the dependence between the features (see below). The PAM was initially developed for identification of gene signatures across multiple subtypes of cancer using gene expression microarray data (Tibshirani et al, 2002). It computes group centroids for each gene, and using a soft thresholding method, it shrinks the centroid towards the overall centroid, eventually eliminating the gene for the prediction if it is not predictive. On the other hand, genes with group centroids away from the overall mean become important predictors for that subtype. However, PAM was designed for a single -omics data and assumes independence among each gene. Furthermore, the method was found to have a bias toward the subgroups with larger sample sizes (Blagus & Lusa, 2010; He & Garcia, 2009). Hence, we proposed several modifications to adapt the method to network-wise analysis, including: (1) the use of weighted group centroids for each feature to account for the varying sample size per group, (2) the use of a new test statistic, 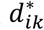, to account for feature-to-feature dependence on the network, and (3) the use of group-specific thresholds in the soft-thresholding method, instead of a uniform threshold (see **Supplementary Information** for details).

Lastly, we carry out pathway enrichment analysis to report biological functions and pathways enriched in the selected edges for each phenotypic group. For the set of selected edges from each phenotypic group, we used the hypergeometric distribution to compute the probability of over-enrichment out of the number of all possible edges given a particular gene function or pathway. The module carries out the test for each phenotypic group, separately for the edges that are up- and down-regulated.

### Modified test statistic for edges

In the PAM, *d_ik_* is defined to be the *t*-statistic for each gene *i*, comparing sample group *k* to the overall centroid:

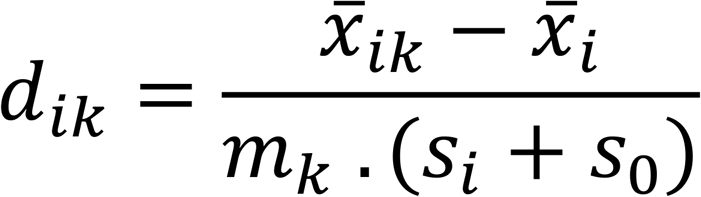

where *x̄_ik_* represents the centroid of gene *i* in phenotypic group *k* and *x̄_i_* represents the overall centroid of gene *i*. The denominator serves as a normalization constant, where 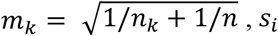 is the pooled within-class standard deviation of gene *i*, and s_4_ is the median value of *S* across all features, a positive constant.

In iOmicsPASS, instead of genes, we use the term *d_ik_* to refer to the test statistic for each edge *i* in phenotypic group *k*. Here, an edge represents an interaction between two molecules, such as a TF protein and mRNA molecule of its target gene. We add an additional term in the test statistic, which is then multiplied by a factor to reward or penalize *d_ik_*, depending on the consistency of co-expression levels in the neighboring edges that share common molecules (nodes). Given a pair of interacting molecules of a particular edge, we define *neighboring edges* as other edges that share one common molecule with it. If the co-expression values of the neighboring edges are mostly up- or down-regulated in a sample group and the edge also has consistent co-expression value, then we reward the edge by adding a term to the test statistics. On the other hand, if the co-expression values of the neighboring edges are not coherent, then the test statistic is penalized (shrunken toward zero).

Specifically, we define 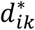 as the new test statistic for edge *i*, comparing phenotypic group *k* to the overall centroid:

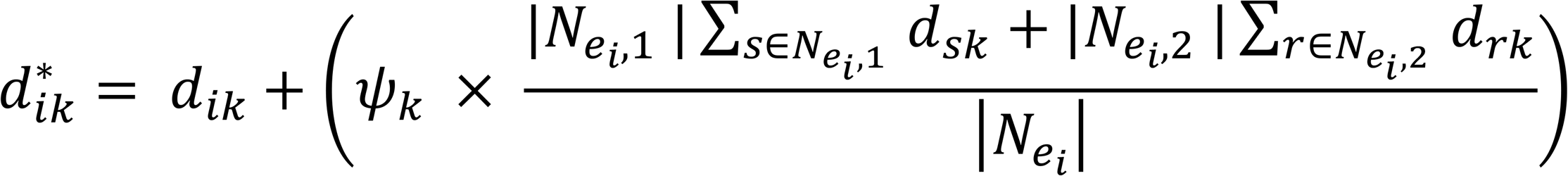

where *N_e_i__* represents the set of neighboring edges for the edge *e_i_*, and |*N_e_i__*| implies the number of edges in the neighborhood set *N_e_i__*. The set, *N_e_i__*, can be further partitioned into two subsets, *N_e_i_,1_* and *N_e_i_,2_*, representing the set of TF and PPI edges, respectively. Here, *ψ_k_* acts as a multiplicative factor for sample group *k* that represents the proportion of agreement between *e_i_* and its neighbouring edges.

### Cross-validation for optimal shrinkage parameter selection

We implemented *K*-fold cross-validation in iOmicsPASS to identify the optimal shrinkage parameter that gives the best classification performance. We split the data split into *K* mutually exclusive parts. Then we use (K-1) folds of data to train the prediction model and use the remaining fold as test data. For a fixed group-specific threshold, ∆*k*, we compute the shrunken centroid 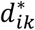

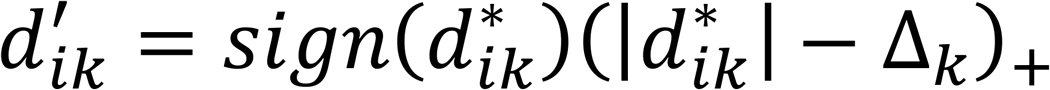

where 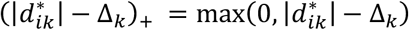. In addition, we set 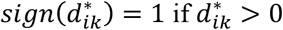, 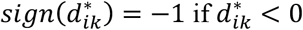 and 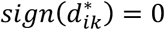, otherwise.

We use the model to predict the membership of test samples and record the misclassification error (or test error). We repeat this process over a grid of increasing values for ∆ starting from 0 to a value sufficiently large so that all the features are reduced towards the overall centroids (i.e. 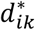 values become zero). This procedure is known as the soft-thresholding method. We repeat the process for each of the *K* folds and average the misclassification errors over the folds to give the overall misclassification error.

### Class probabilities for individual samples

We classify the test samples to the nearest shrunken centroid of each group. Suppose we have a test sample with edge-level co-expression data 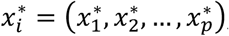 then we compute the discriminant score for group *k* as:

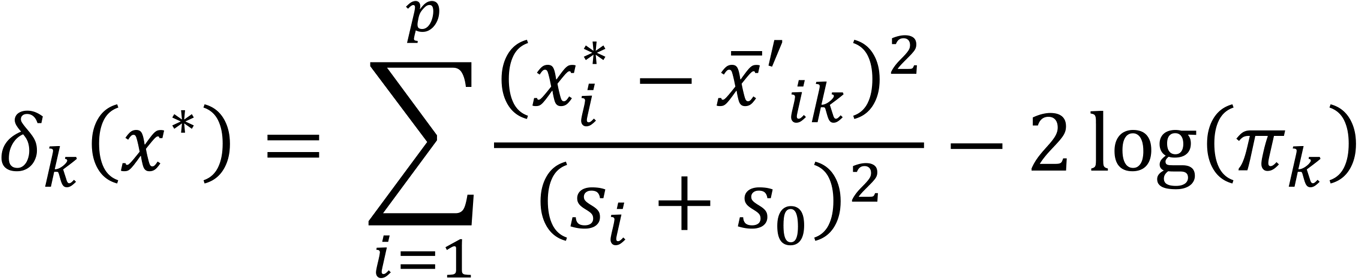

where π_*k*_ is the *k*^th^ class prior probability. We use an equal prior π_*k*_ = 1/*K* Then we assign each test sample to the phenotypic group with the smallest discriminant score using the classification rule: *C*(*x*\*) = *ℓ* where 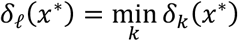. Using the discriminant scores, the estimated probability of sample *x*\* membership to group *k* is computed as:

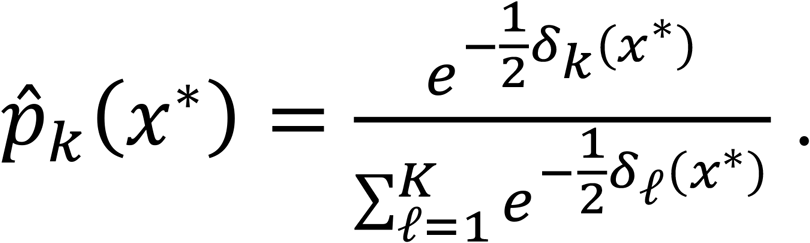

### Simulation Analysis

We randomly selected 1,000 proteins and 5,000 mRNAs where all the proteins are transcription factors and the latter are targets of those transcription factors. A total of 100 sets of protein and mRNA datasets were simulated, assuming that the data was collected from 50 subjects in a disease group and 50 control subjects. We planted the signal, i.e. non-zero centroid, for 100 randomly selected TF proteins and mRNA of all of its interacting partners with stipulated probabilities among the samples of the disease group.

To make the exercise more realistic, we applied a probability called *assay sensitivity* (i.e. *P_AS_*) to determine how likely the signal from the protein is observed in the MS data. In succession from the first, the samples of that protein in the disease group is again generated from Gaussian distribution with mean, *μ*, and standard deviation, *σ* with probability *P_AS_*. Then for the mRNA data, the samples in the disease group were also generated with a probability, *P_mRNA_*, that the TF has real impact on the transcription of the target gene mRNA, proportional to the number of transcription factor targeting it (parent nodes) (see **Figure 2B**). Here, mRNAs with more TF targeting it will have a higher probability of carrying the signal. Simulations were carried out using three different settings: (**A**) signal in the disease group is the strongest with *μ*_Protein_ = *μ*_mRNA_ = 1 and *σ*_Protein_ = *σ*_mRNA_ = 1; (**B**) signal is noisier with *μ*_Protein_ = *μ*_mRNA_ = 0.5 and *σ*_Protein_ = *σ*_mRNA_ = 1; and **(C**) attenuated signal in protein data compared to mRNA data with *μ*_Protein_ = 0.5, *μ*_mRNA_ = 1 and *σ*_Protein_ = *σ*_mRNA_ = 1.

We carried out the simulations as follows:

(1) Randomly select 100 TF proteins to be the true signals.
(2) Identify mRNA targets of the 100 TF proteins, where the signal will be planted.
(3) For each molecule *i* in sample *j*, simulate as follows:

(a) For protein data:

(i) Samples in the disease group are generated with probability *P_AS_*, *X_ij_ ~ N(μ*, *σ*^2^).
(ii) Samples in the control group are generated by *X_ij_ ~ N(*0, *σ*^2^).
(b) For mRNA data:

(i) Samples in the disease group are generated with probability, *P_mRNA_*, *X_ij_ ~ N(μ*, *σ*^2^).
(ii) Samples in the control group are generated by *X_ij_ ~ N(*0, *σ*^2^).
(4) Perform iOmicsPASS analysis on the edge-level data and PAM analysis on the node-level data after concatenating both data together into a single matrix and implement PAM.

Simulations were repeated 100 times and the sensitivities and specificities were averaged across to produce the ROC curve. Both sensitivity and specificity were calculated using an edge-based approach, where the true signal is defined to be an edge that encompasses both the 100 TF proteins and the mRNAs of their target genes (e.g. interacting proteins or mRNAs targets). For PAM, edges were constructed using the set of nodes that are selected and both the protein and mRNA must be selected for the edge to be considered a hit. Sensitivity was calculated using number of true edges selected divided by the number of true edges. Specificity was the number of edges without signal and not selected divided by the number of edges without a signal. Using the set of sensitivities and specificities, the area under the curve (AUC) is then calculated using the trapezoidal rule.

### The Cancer Genome Atlas (TCGA) and The Clinical Proteomic Tumor Analysis Consortium (CPTAC)

Both breast cancer and colorectal cancer data were downloaded using Genomics Data Commons (GDC) data portal from TCGA and proteomics data were downloaded from CPTAC (Edwards et al, 2015). All transcriptomic and copy number variation data were processed using GDC processing pipelines (with GRCh38 as the reference genome).

To standardize the type of gene identifiers used across the different -omics data, all identifiers were mapped to HGNC gene symbol. ENSEMBL identifiers were converted to gene symbols in the mRNA data and gene symbols were already provided in the proteomics data. For CNV data, the chromosomal positions were used to infer the genomic coordinates and a gene symbol was then assigned. If two or more genes overlap at the start and end site, a weighted average of the segment mean values was computed and assigned to each gene. Here, the weights are proportional to the length of the segment that the gene occupies. Then, the average segment mean was computed for each gene in each individual to obtain a single segmental mean value per gene.

For the purpose of our analysis, metastatic samples and normal samples (blood derived and solid tissues) were removed. For subjects with multiple analytes quantified, high correlations between samples were observed and hence, average was taken to yield one sample per individual. Both colon adenocarcinoma (COAD) and rectum adenocarcinoma (READ) were put together and analyzed as colorectal cancer.

### Biological networks and pathway data

We downloaded two types of biological networks: PPI network and TF regulatory network. For the PPI network, the two main sources were iRefIndex (Razick et al, 2008) and BioPlex 2.0 (Huttlin et al, 2017), where we removed the redundancies and compiled into a single database. For the TF regulatory network, we collected the interactions between transcription factor proteins and target genes and put together from the following sources: TRED (Zhao et al, 2005), ITFP (Zheng et al, 2008), ENCODE, and TTRUST (Han et al, 2015). The final network consisted of 14,368 proteins forming 145,906 edges in the PPI network and 1951 TFs and 19,594 target genes forming 284,838 edges in the TF network.

For the pathway data, we used biological pathways from the ConsensusPathDB (CPDB) (Kamburov et al, 2011) and Gene Ontology (GO) (Ashburner et al, 2000). The pathways from the CPDB include data from multiple sources such as KEGG, PharmGKB, SMPDB, HumanCyc, BioCarta, EHMN, Reactome, NetPath, Pathway Interaction Database (PID) and Wikipathways. For GO, we considered the biological processes only. As a result, the final pathway consisted of a total of 14,599 pathways involving 17,248 genes.

All network and pathway files are distributed along with the tool.

### Software Implementation and Computation time

iOmicsPASS is a platform-independent tool, written in C++ language, that uses command line interface. The tool is freely available at Github: https://github.com/cssblab/iOmicsPASS. It is distributed along with a software manual with instructions for installations and example datasets for executing iOmicsPASS. Supplemental *R* codes are provided for the visualization of misclassification errors (cross-validation) and for the choice of appropriate thresholds.

In large-scale data, iOmicsPASS can be time consuming due to multiple high-dimensional -omics datasets. The current total run-time is proportional to the number of samples and the number of interactions constructed from the biological networks, where the latter is more crucial. For example, with ~500,000 edges, the total run time may take up to 10 hours, whereas with ~15,000 edges, the run time is approximately 5 minutes.

For the breast cancer data, a total of 165,296 edges were constructed across 103 samples and the total run time was approximately 4.5 hours. For colorectal cancer, 58,264 edges were constructed across 96 samples and the total run time was about 45 minutes.

## Acknowledgement

This work was supported in part by Institute of Molecular & Cell Biology, ASTAR, and a grant from Singapore Ministry of Education (to HC and KC; MOE2016-T2-1-001) and National Medical Research Council of Singapore (to HC; NMRC-CG-M009). Authors thank Christine Vogel for critical reading and helpful suggestions for the manuscript.

## Author contributions

H.K. and H.C. conceived the project. H.K., K.C. and H.C. developed the algorithm and H.K. and D.F. implemented the software. R.E. led biological interpretation of the data analysis results. H.K. and H.C. wrote the manuscript with input from all authors. H.C. supervised the project.

## Conflict of interest

The authors declare that they have no conflict of interest.

## References

Ashburner M, Ball CA, Blake JA, Botstein D, Butler H, Cherry JM, Davis AP, Dolinski K, Dwight SS, Eppig JT, Harris MA, Hill DP, Issel-Tarver L, Kasarskis A, Lewis S, Matese JC, Richardson JE, Ringwald M, Rubin GM, Sherlock G (2000) Gene ontology: tool for the unification of biology. The Gene Ontology Consortium. Nat Genet 25: 25-29

Becker-Santos DD, Lonergan KM, Gronostajski RM, Lam WL (2017) Nuclear Factor I/B: A Master Regulator of Cell Differentiation with Paradoxical Roles in Cancer. EBioMedicine 22: 2-9

Bersanelli M, Mosca E, Remondini D, Giampieri E, Sala C, Castellani G, Milanesi L (2016) Methods for the integration of multi-omics data: mathematical aspects. BMC Bioinformatics 17 Suppl 2: 15

Blagus R, Lusa L (2010) Class prediction for high-dimensional class-imbalanced data. BMC Bioinformatics 11: 523

Bonnet E, Calzone L, Michoel T (2015) Integrative multi-omics module network inference with Lemon-Tree. PLoS Comput Biol 11: e1003983

Cancer Genome Atlas N (2012) Comprehensive molecular portraits of human breast tumours. Nature 490: 61-70

De Sousa EMF, Wang X, Jansen M, Fessler E, Trinh A, de Rooij LP, de Jong JH, de Boer OJ, van Leersum R, Bijlsma MF, Rodermond H, van der Heijden M, van Noesel CJ, Tuynman JB, Dekker E, Markowetz F, Medema JP, Vermeulen L (2013) Poor-prognosis colon cancer is defined by a molecularly distinct subtype and develops from serrated precursor lesions. Nat Med 19: 614-618

Edwards NJ, Oberti M, Thangudu RR, Cai S, McGarvey PB, Jacob S, Madhavan S, Ketchum KA (2015) The CPTAC Data Portal: A Resource for Cancer Proteomics Research. J Proteome Res 14: 2707-2713

Guinney J, Dienstmann R, Wang X, de Reynies A, Schlicker A, Soneson C, Marisa L, Roepman P, Nyamundanda G, Angelino P, Bot BM, Morris JS, Simon IM, Gerster S, Fessler E, De Sousa EMF, Missiaglia E, Ramay H, Barras D, Homicsko K et al (2015) The consensus molecular subtypes of colorectal cancer. Nat Med 21: 1350-1356

Guiu S, Michiels S, Andre F, Cortes J, Denkert C, Di Leo A, Hennessy BT, Sorlie T, Sotiriou C, Turner N, Van de Vijver M, Viale G, Loi S, Reis-Filho JS (2012) Molecular subclasses of breast cancer: how do we define them? The IMPAKT 2012 Working Group Statement. Ann Oncol 23: 2997-3006

Han H, Shim H, Shin D, Shim JE, Ko Y, Shin J, Kim H, Cho A, Kim E, Lee T, Kim H, Kim K, Yang S, Bae D, Yun A, Kim S, Kim CY, Cho HJ, Kang B, Shin S et al (2015) TRRUST: a reference database of human transcriptional regulatory interactions. Sci Rep 5: 11432

He H, Garcia EA (2009) Learning from imbalanced data. IEEE Transactions on Knowledge and Data Engineering 21: 1263-1284

Huang S, Chaudhary K, Garmire LX (2017) More Is Better: Recent Progress in Multi-Omics Data Integration Methods. Front Genet 8: 84

Huttlin EL, Bruckner RJ, Paulo JA, Cannon JR, Ting L, Baltier K, Colby G, Gebreab F, Gygi MP, Parzen H, Szpyt J, Tam S, Zarraga G, Pontano-Vaites L, Swarup S, White AE, Schweppe DK, Rad R, Erickson BK, Obar RA et al (2017) Architecture of the human interactome defines protein communities and disease networks. Nature 545: 505-509

Ideker T, Ozier O, Schwikowski B, Siegel AF (2002) Discovering regulatory and signalling circuits in molecular interaction networks. Bioinformatics 18 Suppl 1: S233-240

Kamburov A, Pentchev K, Galicka H, Wierling C, Lehrach H, Herwig R (2011) ConsensusPathDB: toward a more complete picture of cell biology. Nucleic Acids Res 39: D712-717

Kim D, Li R, Dudek SM, Ritchie MD (2013) ATHENA: Identifying interactions between different levels of genomic data associated with cancer clinical outcomes using grammatical evolution neural network. BioData Min 6: 23

Kim M, Tagkopoulos I (2018) Data integration and predictive modeling methods for multi-omics datasets. Mol Omics 14: 8-25

Lai MD, Xu J (2007) Ribosomal proteins and colorectal cancer. Curr Genomics 8: 43-49

Laurent JM, Vogel C, Kwon T, Craig SA, Boutz DR, Huse HK, Nozue K, Walia H, Whiteley M, Ronald PC, Marcotte EM (2010) Protein abundances are more conserved than mRNA abundances across diverse taxa. Proteomics 10: 4209-4212

Leroy C, Fialin C, Sirvent A, Simon V, Urbach S, Poncet J, Robert B, Jouin P, Roche S (2009) Quantitative phosphoproteomics reveals a cluster of tyrosine kinases that mediates SRC invasive activity in advanced colon carcinoma cells. Cancer Res 69: 2279-2286

Li M, Stoneking M (2012) A new approach for detecting low-level mutations in next-generation sequence data. Genome Biol 13: R34

Lin D, Zhang J, Li J, Calhoun VD, Deng HW, Wang YP (2013) Group sparse canonical correlation analysis for genomic data integration. BMC Bioinformatics 14: 245

Linnekamp JF, Hooff SRV, Prasetyanti PR, Kandimalla R, Buikhuisen JY, Fessler E, Ramesh P, Lee K, Bochove GGW, de Jong JH, Cameron K, Leersum RV, Rodermond HM, Franitza M, Nurnberg P, Mangiapane LR, Wang X, Clevers H, Vermeulen L, Stassi G et al (2018) Consensus molecular subtypes of colorectal cancer are recapitulated in in vitro and in vivo models. Cell Death Differ 25: 616-633

Lock EF, Hoadley KA, Marron JS, Nobel AB (2013) Joint and Individual Variation Explained (Jive) for Integrated Analysis of Multiple Data Types. Ann Appl Stat 7: 523-542

Meldrum C, Doyle MA, Tothill RW (2011) Next-generation sequencing for cancer diagnostics: a practical perspective. Clin Biochem Rev 32: 177-195

Mertins P, Mani DR, Ruggles KV, Gillette MA, Clauser KR, Wang P, Wang X, Qiao JW, Cao S, Petralia F, Kawaler E, Mundt F, Krug K, Tu Z, Lei JT, Gatza ML, Wilkerson M, Perou CM, Yellapantula V, Huang KL et al (2016) Proteogenomics connects somatic mutations to signalling in breast cancer. Nature 534: 55-62

Meyer M, Briggs AW, Maricic T, Hober B, Hoffner B, Krause J, Weihmann A, Paabo S, Hofreiter M (2008) From micrograms to picograms: quantitative PCR reduces the material demands of high-throughput sequencing. Nucleic Acids Res 36: e5

Mo Q, Wang S, Seshan VE, Olshen AB, Schultz N, Sander C, Powers RS, Ladanyi M, Shen R (2013) Pattern discovery and cancer gene identification in integrated cancer genomic data. Proc Natl Acad Sci U S A 110: 4245-4250

Pandey A, Liu X, Dixon JE, Di Fiore PP, Dixit VM (1996) Direct association between the Ret receptor tyrosine kinase and the Src homology 2-containing adapter protein Grb7. J Biol Chem 271: 10607-10610

Parker JS, Mullins M, Cheang MC, Leung S, Voduc D, Vickery T, Davies S, Fauron C, He X, Hu Z, Quackenbush JF, Stijleman IJ, Palazzo J, Marron JS, Nobel AB, Mardis E, Nielsen TO, Ellis MJ, Perou CM, Bernard PS (2009) Supervised risk predictor of breast cancer based on intrinsic subtypes. J Clin Oncol 27: 1160-1167

Pasquale EB (2010) Eph receptors and ephrins in cancer: bidirectional signalling and beyond. Nat Rev Cancer 10: 165-180

Razick S, Magklaras G, Donaldson IM (2008) iRefIndex: a consolidated protein interaction database with provenance. BMC Bioinformatics 9: 405

Robin JD, Ludlow AT, LaRanger R, Wright WE, Shay JW (2016) Comparison of DNA Quantification Methods for Next Generation Sequencing. Sci Rep 6: 24067

Schwender H (2012) Imputing missing genotypes with weighted k nearest neighbors. J Toxicol Environ Health A 75: 438-446

Takeno S, Sakai T (1991) Involvement of the intestinal microflora in nitrazepam-induced teratogenicity in rats and its relationship to nitroreduction. Teratology 44: 209-214

Tibshirani R, Hastie T, Narasimhan B, Chu G (2002) Diagnosis of multiple cancer types by shrunken centroids of gene expression. Proc Natl Acad Sci U S A 99: 6567-6572

Vaske CJ, Benz SC, Sanborn JZ, Earl D, Szeto C, Zhu J, Haussler D, Stuart JM (2010) Inference of patient-specific pathway activities from multi-dimensional cancer genomics data using PARADIGM. Bioinformatics 26: i237-245

Vinatzer U, Dampier B, Streubel B, Pacher M, Seewald MJ, Stratowa C, Kaserer K, Schreiber M (2005) Expression of HER2 and the coamplified genes GRB7 and MLN64 in human breast cancer: quantitative real-time reverse transcription-PCR as a diagnostic alternative to immunohistochemistry and fluorescence in situ hybridization. Clin Cancer Res 11: 8348-8357

Vu T, Datta PK (2017) Regulation of EMT in Colorectal Cancer: A Culprit in Metastasis. Cancers (Basel) 9

Wei Z, Li H (2007) A Markov random field model for network-based analysis of genomic data. Bioinformatics 23: 1537-1544

Yang X, Han H, De Carvalho DD, Lay FD, Jones PA, Liang G (2014) Gene body methylation can alter gene expression and is a therapeutic target in cancer. Cancer Cell 26: 577-590

Yousef EM, Furrer D, Laperriere DL, Tahir MR, Mader S, Diorio C, Gaboury LA (2017) MCM2: An alternative to Ki-67 for measuring breast cancer cell proliferation. Mod Pathol 30: 682-697

Yuan Y, Savage RS, Markowetz F (2011) Patient-specific data fusion defines prognostic cancer subtypes. PLoS Comput Biol 7: e1002227

Zahnow CA (2009) CCAAT/enhancer-binding protein beta: its role in breast cancer and associations with receptor tyrosine kinases. Expert Rev Mol Med 11: e12

Zhang B, Wang J, Wang X, Zhu J, Liu Q, Shi Z, Chambers MC, Zimmerman LJ, Shaddox KF, Kim S, Davies SR, Wang S, Wang P, Kinsinger CR, Rivers RC, Rodriguez H, Townsend RR, Ellis MJ, Carr SA, Tabb DL et al (2014) Proteogenomic characterization of human colon and rectal cancer. Nature 513: 382-387

Zhao F, Xuan Z, Liu L, Zhang MQ (2005) TRED: a Transcriptional Regulatory Element Database and a platform for in silico gene regulation studies. Nucleic Acids Res 33: D103-107

Zheng G, Tu K, Yang Q, Xiong Y, Wei C, Xie L, Zhu Y, Li Y (2008) ITFP: an integrated platform of mammalian transcription factors. Bioinformatics 24: 2416-2417

